# Chromogranin A plays a crucial role in the age-related development of insulin resistance and hypertension

**DOI:** 10.1101/2021.11.08.467835

**Authors:** Matthew A. Liu, Suborno Jati, Kechun Tang, Geert van den Bogaart, Gourisankar Ghosh, Sushil K. Mahata

**Author notes:** The authors have nothing to declare. Address correspondences to: Sushil K. Mahata, Ph.D. Metabolic Physiology & Ultrastructural Biology Laboratory, Department of Medicine, University of California, San Diego (0732), 9500 Gilman Drive, La Jolla, CA 92093-0732.

## Abstract

Aging is associated with the development of metabolic disorders, including insulin resistance and hypertension. Young mice that are negative for the neuroendocrine prohormone Chromogranin A (CgA knockout, CgA-KO) display two opposite aging phenotypes: hypertension but heightened insulin sensitivity. We determined these phenotypes in aging CgA mice. In comparison, aging wild-type (WT) mice gradually lost glucose tolerance and insulin sensitivity. Moreover, while aging WT mice had increased inflammation with higher plasma TNF-α, IFN-γ and CCL2 and increased mitochondrial fission, these phenotypes were the opposite in aging CgA-KO mice. CgA-KO mice also showed increased expression of mitochondrial and nuclear-encoded complex I genes, implying that they were healthier than WT mice. Most intriguingly, the hypertension in CgA-KO mice was spontaneously reversed with aging. Supplementation of CgA-KO mice with pancreastatin, a hyperglycemic peptide produced from CgA by proteolysis, increased both blood glucose levels and blood pressure, implicating hyperglycemia, and hypertension. We conclude that age-related insulin resistance and hypertension are caused by CgA.

## INTRODUCTION

Insulin resistance is associated with the development of age-related diseases like type 2 diabetes, hypertension, coronary heart disease, stroke, and cancer (1). Similarly, several studies have found an association between aging and a decline in insulin sensitivity in the general population (1; 2). In mice, the association of aging with increased insulin resistance has also been reported (3). The following mechanisms have been implicated in this age-related decrease in insulin sensitivity: (i) defects in insulin signaling (4), (ii) a decrease in insulin-stimulated whole-body glucose oxidation (4), (iii) a reduction in the β-cell response to glucose (4), (iv), impaired insulin-mediated glucose uptake (5), and (v) inability to suppress hepatic glucose production (5). Despite having a general understanding of these mechanisms behind aging and insulin resistance, their exact control mechanisms are less clear.

Evidence suggests that aging is under hormonal control. Mutations in the *Prop-1* (Ames dwarf mice carrying the *Prop-1* mutation) and *Pit-1* (Snell dwarf mice carrying the *Pit-1* mutation) (6) genes cause a life-long combined hormonal deficiency in growth hormone (GH), thyroid stimulating hormone, and prolactin, and increase the longevity of the mice. Other studies found similar results, as GH receptor disrupted GHR-KO mice with profound GH resistance, and GH releasing hormone disrupted (GHRH-KO) mice with isolated GH deficiency (7), also had increased longevity compared to WT mice. Age-related insulin resistance might also be under hormonal control, since the increased longevity in Ames dwarf, Snell dwarf, GHR-KO, and GHRH-KO mice were strongly correlated with enhanced insulin sensitivity (8).

This study examined the role of another hormone in age-related insulin resistance: Chromogranin A (CgA). CgA is a ~49 kDa proprotein, which is the precursor to several biologically active peptides with counterregulatory functions, particularly pancreastatin (PST: hCgA_250-301_) and catestatin (CST: hCgA_352-372_). While PST is a hyperglycemic, prodiabetic (9), pro-inflammatory and pro-obesogenic (10; 11) peptide, CST is an antidiabetic (12), anti-inflammatory (12; 13), anti-obesogenic (14) and anti-hypertensive (15) peptide. Although CgA is overexpressed in hypertensive humans and rodents, the CST peptide is low in hypertensive subjects as well as in offspring of hypertensive parents (16). Furthermore, we previously reported that while CgA knockout (CgA-KO) mice are hypertensive owing to the lack of CST (15), they are also insulin-sensitive owing to the lack of PST, even after 4 months on a high-fat diet (HFD: 60% calorie from fat) (10; 11).

Since both the human and rodent studies indicate that aging correlates with insulin resistance and hypertension, we determined how these phenotypes were affected in aging CgA-KO mice. The data demonstrate increased insulin sensitivity in aging CgA-KO mice, which is accompanied by a spontaneous reversal of the hypertension phenotype. These could be inversed by injection of PST, indicating that age-related insulin resistance and hypertension are caused by CgA.

## RESEARCH DESIGN AND METHODS

### Animals and Diets

Male WT and CgA-KO (0.5 to 2 yrs old) were in C57BL/6 background. Since CgA is especially overexpressed in male patients with hypertension (17), we used only male mice in this study. Mice were kept in a 12 hr dark/light cycle and fed a normal chow diet (NCD: 13.5% calorie from fat; LabDiet 5001, TX). Animals were age-matched, and randomly assigned for each experiment. Control and experimental groups were blinded. All studies with mice were approved by the UCSD and Veteran Affairs San Diego Institutional Animal Care and Use Committees and conform to relevant National Institutes of Health guidelines.

### Glucose tolerance test (GTT), glucose-stimulated insulin secretion (GSIS), and insulin tolerance test (ITT)

For GTT, glucose (1 mg/g body weight) was injected intraperitoneally (time zero) after an 8-hr fast. Tail-vein glucose levels were measured at 0, 15, 30, 60, 90, and 120 min. For GSIS, blood was collected from the tail-vein at 0 and 10 min and the plasma was used for an insulin assay. For insulin tolerance tests (ITT), insulin (0.4 mU/g body weight) was injected intraperitoneally, and blood glucose levels were measured at the same time points as for GTT.

### Protein analysis by immunoblotting

Liver pieces were homogenized in a buffer (50 mM Tris–Cl pH 8.0, 150 mM NaCl, 1% Triton X-100, 0.5% sodium deoxycholate, 0.1% SDS, 50 mM DTT, 5% glycerol, 5 mM NaF, 2 mM Na3VO4, 50 mM PMSF, 1 mM EDTA, Protease Inhibitor Cocktail from Sigma Aldrich) as previously described (17). Liver homogenates were subjected to SDS-PAGE and immunoblotted with antibodies directed against Phospho-Ser473-AKT (1:2000; catalog #9271S) and total AKT (1:4000; catalog #4685S), phospho-Ser9-GSK-3β (1:2000; catalog #9322S) and total GSK-3β (1:4000; catalog #9832S), phospho-Thr180/Tyr182-p38 (1:2000; catalog #9211S) and total p38 (1:5000; catalog #9212S) as well as phospho-Thr202/Tyr204-ERK1/2 (1:1000; catalog #4377S) and total ERK1/2 (1:3000; catalog #4695S). All these primary antibodies were purchased from Cell Signaling Technology (Danvers, MA). Anti-rabbit IgG-HRP conjugate (A6154; 1:8000) and anti-mouse IgG-HRP conjugate (A9044; 1:8000) were purchased from Sigma-Aldrich (St. Louis, MO).

### Electron microscopy

To displace blood and wash off tissues before fixation, mice were cannulated through the apex of the heart and perfused with a calcium and magnesium free buffer composed of DPBS (Life Technologies Inc), 10 mM HEPES, 0. 2 mM EGTA, 0.2% BSA, 5 mM glucose and KCl concentration adjusted 9.46 mM (to arrest the heart in diastole) as described previously (18). This was followed by perfusion fixation with freshly prepared fixative containing 2.5% glutaraldehyde, 2% paraformaldehyde in 0.15 M cacodylate buffer, and postfixed in 1% OsO4 in 0.1 M cacodylate buffer for 1 hour on ice. After perfusion, small pieces of the liver were immersed in the above fixative for 12-16 hrs. The tissues were stained *en bloc* with 2-3% uranyl acetate for 1 hour on ice. The tissues were dehydrated in graded series of ethanol (20-100%) on ice followed by one wash with 100% ethanol and two washes with acetone (15 min each) and embedding with Durcupan. Sections were cut at 50 to 60 nm on a Leica UCT ultramicrotome and picked up on Formvar and carbon-coated copper grids. Sections were stained with 2% uranyl acetate for 5 minutes and Sato’s lead stain for 1 minute. Grids were viewed using a JEOL JEM1400-plus TEM (JEOL, Peabody, MA) and photographed using a Gatan OneView digital camera with 4k x 4k resolution (Gatan, Pleasanton, CA).

### Real Time PCR

Total RNA from liver tissue was isolated using RNeasy Mini Kit and reverse-transcribed using a qScript cDNA synthesis kit. cDNA samples were amplified using PERFECTA SYBR FASTMIX L-ROX 1250 and analyzed on an Applied Biosystems 7500 Fast Real-Time PCR system. All PCRs were normalized to *Rplp0* (Ribosomal protein, large, P0), and relative expression levels were determined by the ΔΔ*C_t_* method.

### Measurement of cytokines

Plasma cytokines or supernatant of cultured BMDMs (20 μl) were measured using U-PLEX mouse cytokine assay kit (Meso Scale Diagnostics, Rockville, MD) via the manufacturer’s protocol.

### Tail-cuff measurement of blood pressure

Systolic blood pressure (SBP) was measured using the mouse and rat tail cuff blood pressure (MRBP) System (IITC Life Sciences Inc. Woodland Hills, CA). Mice were restrained in plexiglass tubes and heated to 34°C for 10-15 min in individual warming chambers prior to BP measurement. The tails were placed inside inflatable cuffs with a photoelectric sensor that measured tail pulses. The SBP was measured over 6 separate days with an average of two values per day.

### Statistics

Statistics were performed with PRISM 8 (version 8.4.3) software (San Diego, CA). Data were analyzed using unpaired two-tailed Student’s t-test for comparison of two groups or 1-way/2-way/3-way analysis of variance (ANOVA) for more than two groups followed by Tukey’s post hoc test if appropriate. All data are presented as mean ± SEM. Significance was assumed when p<0.05.

## Results

### Aging CgA-KO mice display improved insulin sensitivity

It is well documented that aging is associated with declines in glucose tolerance and insulin sensitivity in both mice (3) and humans (19). We have reported previously that young (~4-6 months old) CgA-KO mice on both a normal chow diet (NCD) and a high fat diet display heightened insulin sensitivity (10; 11). Therefore, we asked the question whether CgA-KO mice could maintain improved insulin sensitivity with aging. As expected (10), the glucose tolerance test (GTT) revealed a gradual decrease in glucose tolerance with aging in WT mice (Fig. 1A&B). Likewise, glucose-stimulated insulin secretion (GSIS) was reduced in 1 and 2 yr old WT mice (Fig. 1C). In contrast, improved glucose tolerance and GSIS were maintained in aging CgA-KO mice (Fig. 1A-C). The insulin tolerance test (ITT) showed progressive deterioration of insulin sensitivity in aging (0.5 yr to 2 yr) WT mice (Fig. 1D&E). In contrast, aging CgA-KO mice showed progressive improvements in insulin sensitivity (0.5 to 2 yr) (Fig. 1D&E).

**Fig. 1.**
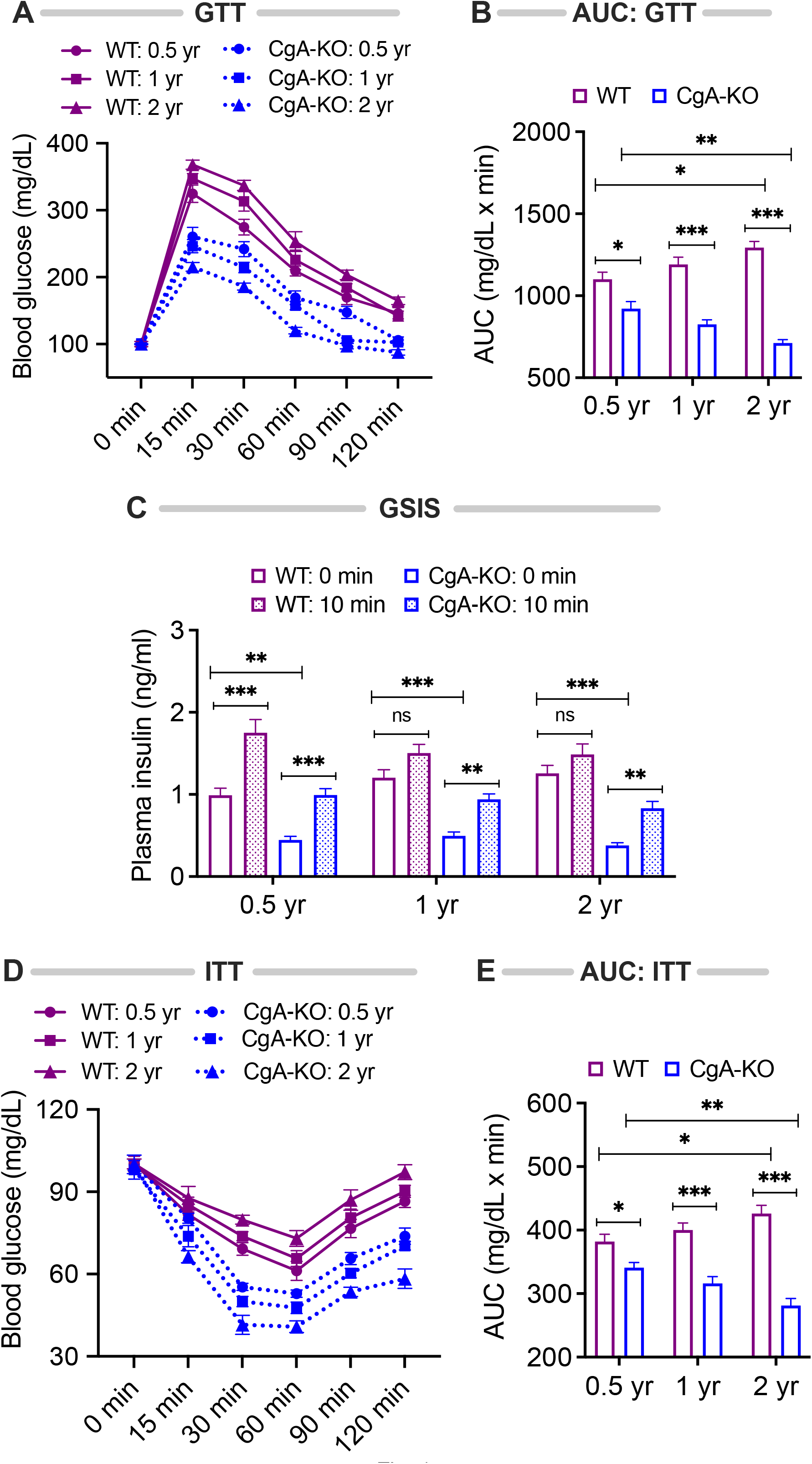
Aging CgA-KO mice maintain and improve glucose tolerance and insulin sensitivity. (A) Glucose tolerance test (GTT, n=7) with 8h fasting mice and (B) the corresponding area under the curve (AUC). 2-way ANOVA. Interaction: p<0.001; Time: p<0.001; Genotype and age: p<0.001. (C) Glucose-stimulated insulin secretion (GSIS) with 8h fasting mice. Data were analyzed by 3-way ANOVA: Age: ns; Genotype; p<0.001; Treatment: p<0.001; Age x Genotype: ns; Age x treatment: p<0.05; Genotype x Treatment: ns; Age x Genotype x Treatment: ns. (D) Insulin tolerance test (ITT, n=7) on 8h fasting mice and (E) the corresponding AUC. 2-way ANOVA. Interaction: p<0.001; Time: p<0.001; Genotype and age: p<0.001.

### Aging CgA-KO mice maintain insulin signaling pathways

Insulin promotes glucose uptake by signaling via the serine/threonine kinase AKT (20). While intraperitoneal injection with insulin did not result in increased phosphorylation of AKT (S473) in the liver of 2 yr old WT mice, it did so in 2 yr old CgA-KO mice (Fig. 2A&B). Another important function of insulin is to promote glycogenesis through the activation of glycogen synthase kinase (GSK)-3β by AKT (20). Administration of insulin also did not result in phosphorylation of GSK-3β (S9) in the liver of WT mice, but it did in 2 yr old CgA-KO mice (Fig. 2A&C). The mitogen-activated protein kinase (MAPK) pathway constitutes a second essential branch of insulin signaling, which is responsible for cell growth, differentiation, and survival (21). Insulin-stimulated phosphorylation of MAPK components p38 (T180/Y182) and ERK1/2 (T202/Y204) was compromised in the liver of aging WT mice (2 yr old), but not affected in aging CgA-KO mice (Fig. 2A, D&E). These findings confirm the maintenance of insulin sensitivity in aging CgA-KO mice.

**Fig. 2.**
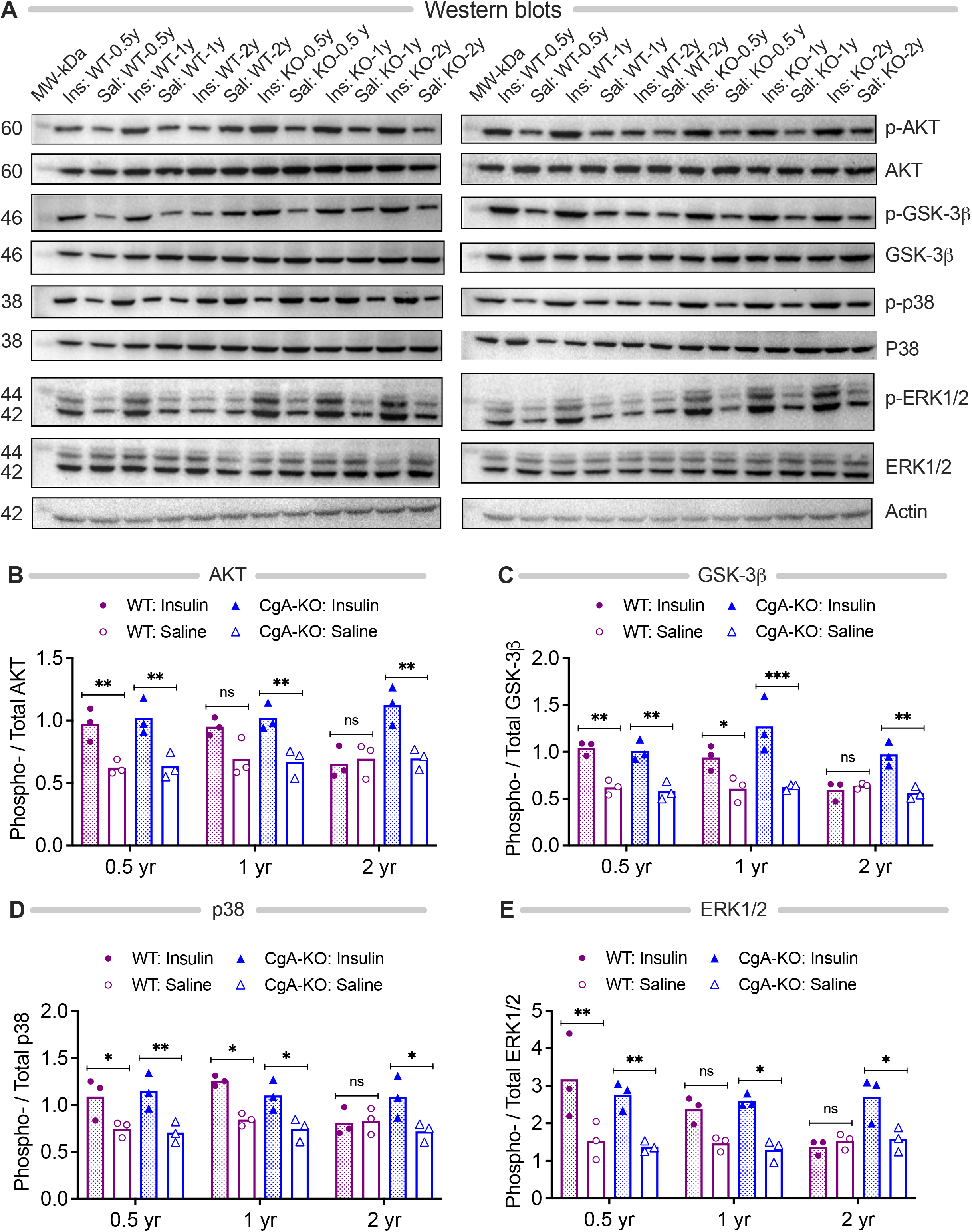
Aging CgA-KO mice maintain insulin signaling pathways. (A) Western blots of phosphorylated-AKT (S473), total AKT, phosphorylated GSK-3β (S9), total GSK-3β, phosphorylated p38 (T180/Y182), total p38, phosphorylated ERK1/2 (T202/Y204), and total ERK1/2 in the livers of 2 sets of aging WT and CgA-KO mice. (B-E) Densitometric analyses of the Western blots by 3-way ANOVA (n=3). (B) AKT (phosphorylated/total AKT). Age: ns; Genotype: p<0.01; Treatment: p<0.001; Age x Genotype: p<0.05; Age x Treatment: ns; Genotype x Treatment: p<0.01; Age x Genotype x Treatment; p<0.05. (C) GSK-3β (phosphorylated/total GSK-3β. 3-way ANOVA: Age: p<0.01; Genotype: ns; Treatment: p<0.001; Age x Genotype: ns; Age x Treatment: p<0.05; Genotype x Treatment: p<0.05; Age x Genotype x Treatment: ns. (D) p38 (phosphorylated/total p38). Age: ns; Genotype: ns; Treatment: p<0.001; Age x Genotype: ns; Age x Treatment: ns; Genotype x Treatment: p<0.05; Age x Genotype x Treatment: p<0.05. (E) ERK1/2 (phosphorylated/total ERK). Age: ns; Genotype: ns; Treatment: p<0.001; Age x Genotype: ns; Age x Treatment: ns; Age x Genotype x Treatment: ns.

### Suppression of hepatic glucose production in aging CgA-KO mice

One of the main functions of insulin is to suppress hepatic glucose production (HGP), which is known to be compromised in the elderly population (2). Consistent with literature (2), our data show increased expression of gluconeogenic genes *Pck1* (phosphoenolpyruvate carboxykinase 1) and *G6pc* (glucose-6-phosphatse) in the liver of aging (1 and 2 yr old) WT mice (Fig. 3A&B). In contrast, this increase was not observed in aging CgA-KO mice (Fig. 3A&B). These data indicate that the elevated insulin sensitivity in CgA-KO is at least partly caused by suppression of HGP.

**Fig. 3.**
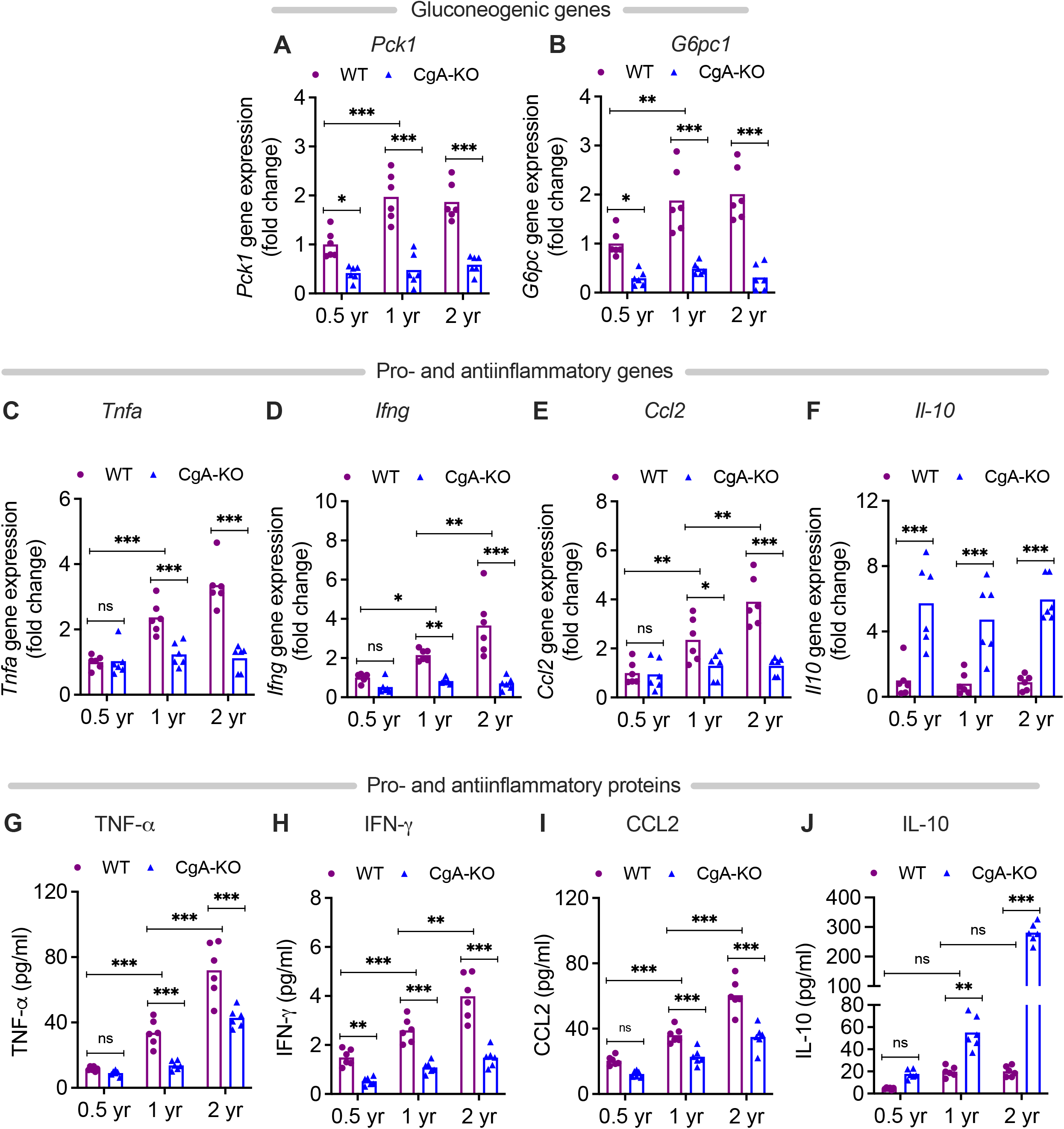
Lower expression of gluconeogenic genes in aging CgA-KO mice. 2-way ANOVA was used to analyze the following gene expression data in the liver of aging WT and CgA-KO mice (n=6): (A) Expression of *Pck1* gene. Interaction: p<0.01; Age: p<0.001; Genotype: p<0.001. (B) Expression of *G6pc* gene. Interaction: p<0.05; Age: p<0.001; Genotype: p<0.01. **Less age-related inflammation in CgA-KO mice.** (C-F) 2-way ANOVA was used to analyze the following gene expression data in the liver of aging WT and CgA-KO mice (n=6): (C) *Tnfa.* Interaction: p<0.001; Age: p<0.001; Genotype: p<0.001. (D) *Ifng.* Interaction: p<0.001; Age: p<0.001; Genotype: p<0.001. (E) *Ccl2*. Interaction: p<0.001; Age: p<0.001; Genotype: p<0.001. (F) *Il10*. Interaction: ns; Age: p<0.001; Genotype: ns. (G-J) 2-way ANOVA was used to analyze the following protein levels in plasma of aging WT and CgA-KO mice (n=6): (G) TNFα. Interaction: p<0.01; Age: p<0.001; Genotype: p<0.001. (H) IFNγ. Interaction: p<0.01; Age: p<0.001; Genotype: p<0.001. (I) CCL2. Interaction: p<0.01; Age: p<0.001; Genotype: p<0.001. (J) IL10. Interaction: p<0.001; Age: p<0.001; Genotype: p<0.001.

### CgA-KO mice maintain an anti-inflammatory status during aging

A low-grade pro-inflammatory state is a hallmark of aging, characterized by the elevation of pro-inflammatory cytokines in the liver as well as in the circulation (22). Consistent with literature (22), the liver of aging (1 and 2 yr old) mice had increased expression of proinflammatory genes *Tnfa* (tumor necrosis factor alpha), *Ifng* (interferon gamma) and *Ccl2* (chemokine C-C motif ligand 2) (Fig. 3C-E). Similarly, the levels in the plasma of the corresponding proteins TNFα, IFNγ, and CCL2 were also elevated (Fig. 3G-I). The expression of anti-inflammatory gene *Il10* (interleukin 10) (Fig. 3F) and protein IL10 (Fig. 3J) were not affected in aging WT mice. Compared to WT mice, the opposite phenotype was observed in aging CgA-KO mice: lower hepatic expression of proinflammatory genes *Tnfa, Ifng* and *Ccl2* (Fig. 3C-E) and higher expression of anti-inflammatory gene *Il10* (Fig. 3F). Similar results were detected in plasma protein levels (Fig. 3G-J). These results show that whereas WT mice display increased inflammation upon aging, this is not present in CgA-KO mice.

### Increased mitochondrial fusion in aging CgA-KO mice

Mitochondria frequently undergo coordinated cycles of fusion (elongation, more efficient respiration) and fission (shortened, less efficient), which are called mitochondrial dynamics, to meet specific cellular requirements (23), but these dynamics are compromised with age (24). Ultrastructural studies revealed increased mitochondrial fusion (by 1.9-fold) in liver cells of aging CgA-KO mice compared to increased fission (by 2.2-fold) in aging WT mice (Fig. 4A-F). In line with this, expression of genes involved in mitochondrial fission *Drp1* (dynamin 1-like) and *Fis1* (fission, mitochondrial 1) were increased in the liver of 2 yr old WT mice, whereas the expression of genes involved in mitochondrial fusion *Mfn1* (mitofusin 1) and *Opa1* (mitochondrial dynamin like GTPase) were increased in aging CgA-KO mice (1 and 2 yr old) (Fig. 4G-J). These experiments show that mitochondrial remodeling is less affected in aging CgA-KO mice.

**Fig. 4.**
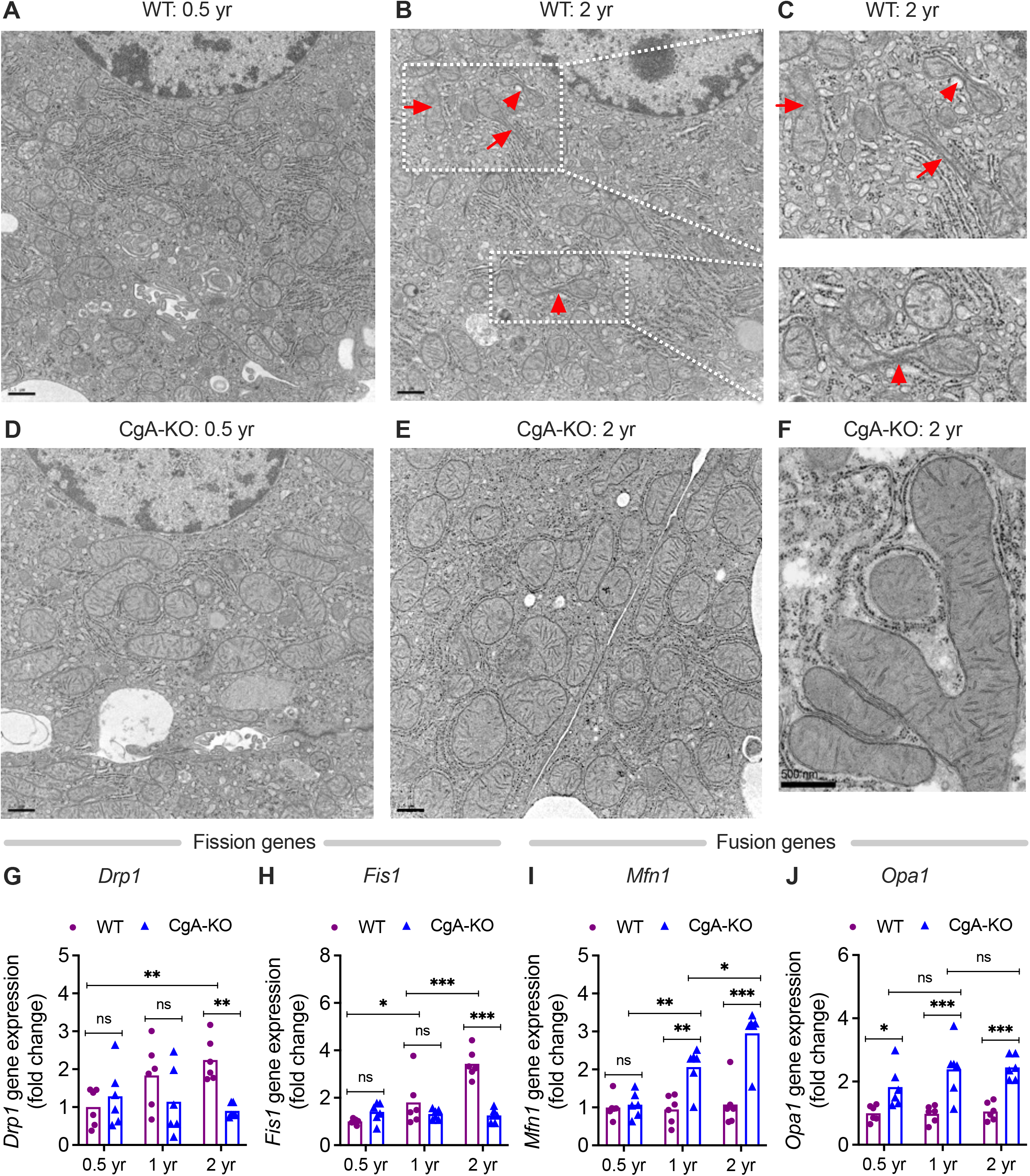
Increased mitochondrial fusion in aging CgA-KO mice. Electron micrographs of liver sections: (A-C) Mitochondrial fission in aging WT mice. (D-F) Mitochondrial fusion in aging CgA-KO mice. Scale bars: 0.5 μm (A, B, D & E). **Expression of mitochondrial fission genes in the liver of WT and CgA-KO mice:** (G) *Drp1.* 2-way ANOVA: Interaction: p<0.05; Age: ns; Genotype: p<0.05. (H) *Fis1.* 2-way ANOVA: Interaction: p<0.001; Age: p<0.001; Genotype: p<0.001. **Expression of mitochondrial fusion genes:** (I) *Mfn1.* 2-way ANOVA: Interaction: p<0.001; Age: p<0.001; Genotype: p<0.001. (J) *Opa1.* 2-way ANOVA: Interaction: ns; Age: ns; Genotype: p<0.001.

### CgA-KO mice exhibit increased mitochondrial biogenesis

Aging is associated with decreases in mitochondrial density, a result of decreased mitochondrial biogenesis (25). *Sirt1* (sirtuin 1) (26), *Sirt6* (sirtuin 6) (27), and *Pgc1α* (peroxisome proliferative activated receptor, gamma, coactivator 1 alpha) (28) are involved in mitochondrial biogenesis and their expression decreases with aging. Although the density of mitochondria was decreased (by 19.8%) in the liver of aging WT mice (Fig. 4A&B), there were no significant changes in the expression of these genes (Fig. 5A-C). In contrast, aging CgA-KO mice showed an increased (by 12.6%) mitochondrial density (Fig. 4D&E) and an increased expression of the *Sirt1, Sirt6,* and *Pgc1a* genes (Fig. 5A-C).

**Fig. 5.**
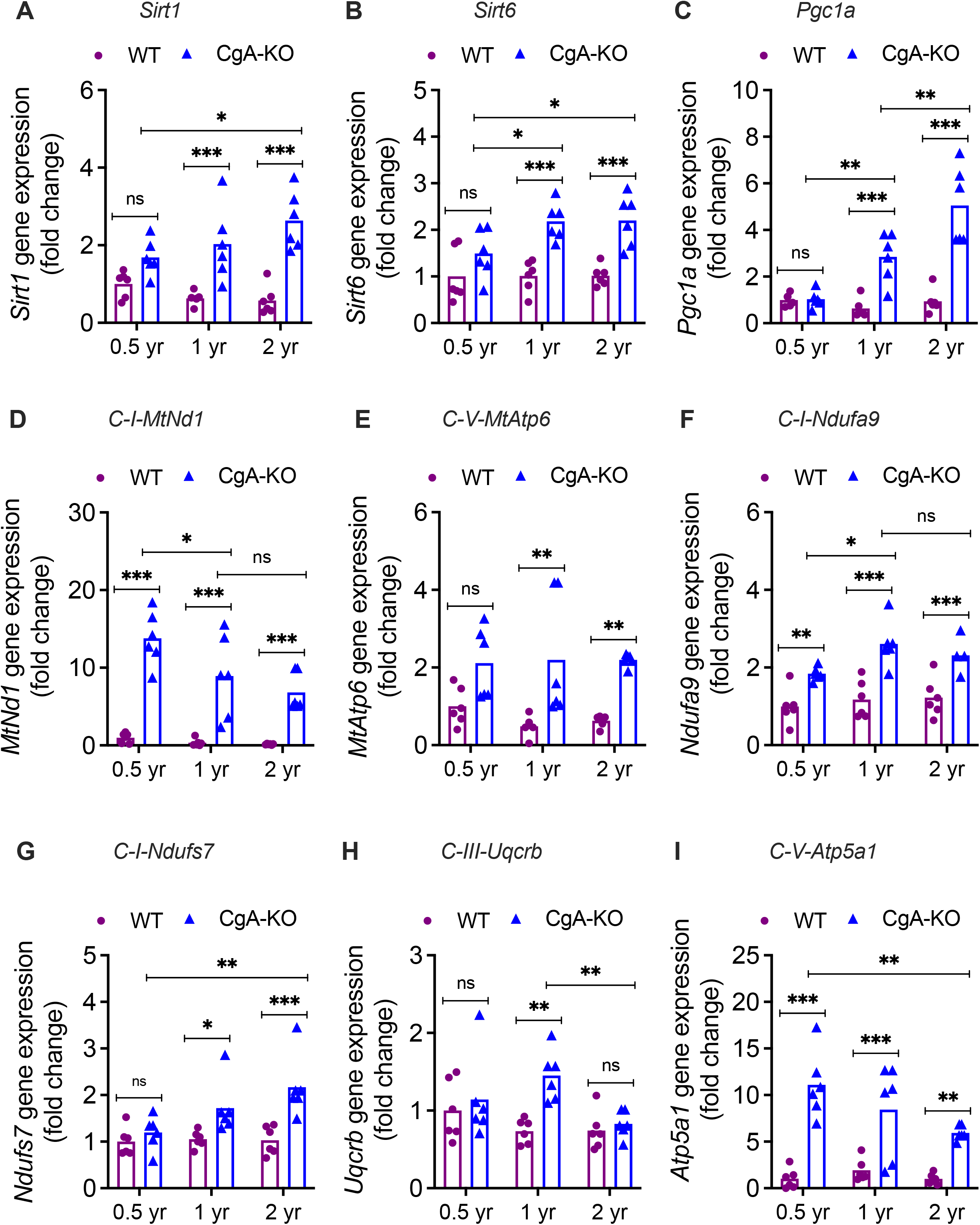
Increased expression of mitochondrial genes in CgA-KO mice. Expression of genes involved in mitochondrial biogenesis in the liver of WT and CgA-KO mice. 2-way ANOVA was used to analyze the following genes (n=6): (A) *Sirt1.* Interaction: p<0.05; Genotype: p<0.001; Age: ns. (B) *Sirt6.* Interaction: ns; Genotype: p<0.001; Age: ns. (C) *Pgc1a.* Interaction: p<0.001; Genotype: p<0.001; Age: p<0.001. **Expression of mitochondrially encoded genes:** (D) *MtNd1.* Interaction: p<0.05; Genotype: p<0.001; Age: p<0.01. (E) *MtAtp6.* Interaction: ns; Genotype: p<0.001; Age: ns. **Expression of nuclear-encoded genes:** (F) *Ndufa9.* Interaction: ns; Genotype: p<0.001; Age: p<0.05. (G) *Ndufs7.* Interaction: p<0.05; Genotype: p<0.001; Age: p<0.05 (H) *Uqcrb.* Interaction: ns; Genotype: p<0.01; Age: ns. (I) Atp5a1. Interaction: ns; Genotype: p<0.001; Age: ns.

### Differential expression of mitochondrial genes in WT and CgA-KO mice

Age-related mitochondrial deterioration (reduced mitochondrial DNA volume, integrity, and functionality) is associated with a decline in mitochondrial function (29). Therefore, we determined expression of mitochondria-encoded genes in aging WT and CgA-KO mice. While the expression of component of Complex I *MtNd1* (mitochondrially encoded NADH dehydrogenase 1) and Complex V *MtAtp6* (mitochondrially encoded ATP synthase 6) decreased in aging WT mice, the expression of those two genes remained consistently high in aging CgA-KO mice compared to WT mice (Fig. 5D&E).

Nuclear-encoded proteins also play crucial roles in optimal functioning of mitochondria (30). Therefore, we also evaluated the expression of nuclear-encoded genes coding for mitochondrial components. The expression of Complex I *Ndufa9* (NADH:ubiquinone oxidoreductase subunit A9) and Complex I *Ndufs7* (NADH:ubiquinone oxidoreductase core subunit S7) were higher in aging CgA-KO mice compared to aging WT mice (Fig. 5F&G). Increased expression of Complex III *Uqcrb* (ubiquinol-cytochrome c reductase binding protein) was evident in 1 yr old CgA-KO mice, but it declined in 2 yr old CgA-KO mice (Fig. 5H). Finally, although CgA-KO displayed increased expression of Complex V *Atp5a1* (ATP synthase, H+ transporting, mitochondrial F1 complex, alpha subunit 1) from 0.5 yr to 2 yr compared to WT mice, there was significant decrease in expression of this gene between 0.5 and 2 yr CgA-KO mice (Fig. 5I). These data indicate that the biogenesis of mitochondria is altered in CgA-KO mice.

### Spontaneous reversal of hypertension in aging CgA-KO mice

Older adults account for the bulk of hypertension-related morbidity and mortality (31). Therefore, we evaluated whether aging increased the blood pressure in mice. Consistent with human data (31), we found a gradual increase in systolic blood pressure (SBP) with aging in WT mice (Fig. 6A). In comparison, while the SBP was increased in young (0.5 yr) CgA-KO mice as we reported previously (15), it was spontaneously reversed in 2 yr old CgA-KO mice compared to their 1 yr old counterparts (Fig. 6A).

**Fig. 6.**
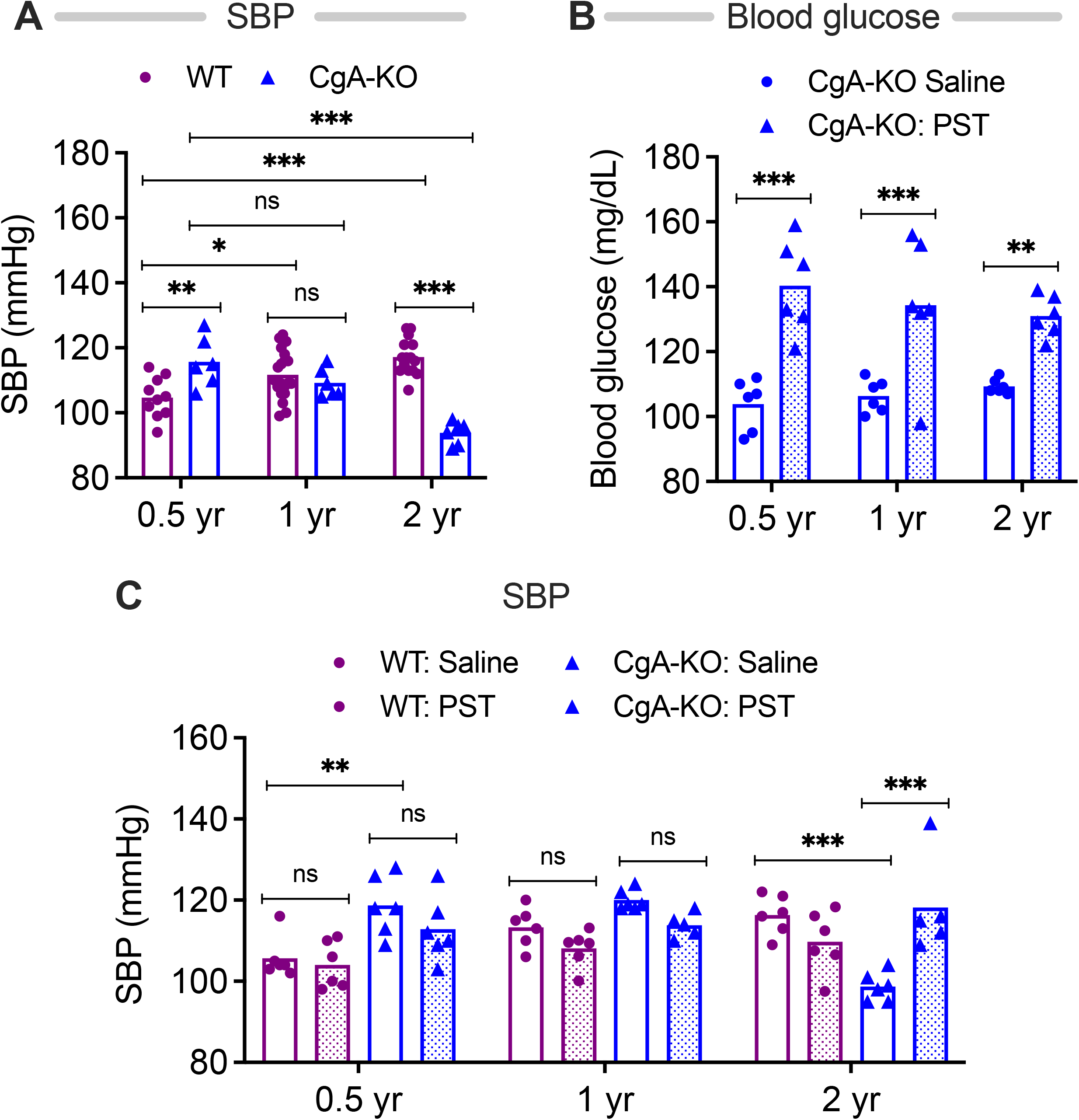
Spontaneous reversal of hypertension in aging CgA-KO mice. (A) Systolic blood pressure (SBP) in aging WT and CgA-KO mice. 2-way ANOVA (n=7-10): Interaction: p<0.001; Genotype: p<0.01; Age: p<0.05. PST increases blood glucose in CgA-KO mice. (B) Blood glucose in PST (2 μg/g BW for 30 days intraperitoneal injection)-treated mice. 2-way ANOVA: Interaction: ns; Age: ns; Genotype & Treatment: p<0.001. PST increases SBP in aging CgA-KO mice. (C) SBP in PST-treated mice. 3-way ANOVA: Age: ns; Genotype: p<0.01; Treatment: ns; Age x Genotype: p<0.001; Age x Treatment: p<0.01; Genotype x Treatment; p<0.05; Age x Genotype x Treatment: p<0.001.

### PST increases blood glucose and SBP in aging CgA-KO mice

One of the reasons for the spontaneous reversal of hypertension in 2 yr old CgA-KO mice could be their increased insulin-sensitivity. PST has been reported to increase blood glucose levels in rodents by activating gluconeogenesis and glycogenolysis (10; 32). Since CgA-KO mice lack PST, we reasoned that supplementation of CgA-KO mice with PST would increase blood glucose levels. Indeed, intraperitoneal administration of PST (2 μg/g body weight/day for one month) increased blood glucose level in 0.5, 1, and 2, yr old CgA-KO mice (Fig. 6B). The increased blood glucose levels upon administration of PST were associated with an increase of SBP in 2 yr old CgA-KO mice (Fig. 6C), suggesting that elevated blood glucose contributes to the hypertension.

## Discussion

The exact mechanisms of aging are incompletely understood. In the current work, we show a role for CgA in aging-related metabolic symptoms. Like humans (31), blood pressure increases in mice as they age, and insulin sensitivity/glucose tolerance decreases. Contrastingly, CgA-KO mice show the inverse with aging. While they initially present with hypertension, this spontaneously reverts between 1-2 years of age. Their insulin sensitivity also improves with age, suggesting that the associated lowering of blood glucose levels might help the switch from a hypertensive to a normotensive phenotype. While the mechanisms of how CgA affects insulin sensitivity during aging are still incompletely understood, we show that the absence of CgA is linked to a decrease in inflammation and more mitochondrial fusion and biogenesis. Both these factors are known to improve insulin sensitivity (33).

The phenotypic changes are not as drastic until around 1 year of age. Therefore, there must be a slow and/or cumulative change that only presents itself phenotypically after a certain period of threshold potential is reached. Moreover, the improved health conditions of CgA-KO mice, as evidenced by progressive insulin sensitivity, a better inflammatory state and mitochondrial health, suggest these mice will live longer beyond the normal lifespan of WT.

Several bioactive peptides are generated from CgA, of which two peptides, CST and PST, display opposite functions (34). We previously showed that CST-KO mice have hypertension (15). These mice also showed hyperinflammation, insulin resistance, high blood glucose levels and poor mitochondrial health. In contrast, PST is known to be hyperglycemic and diabetic (9; 10). Here we show CgA-KO mice become hypertensive and hyperglycemic when injected with the PST peptide, indicating the important role that PST plays in aging-related metabolic symptoms.

### Insulin resistance, impaired glucose tolerance and longevity

Aging is also associated with impaired insulin-mediated glucose uptake, and an inability to suppress hepatic glucose output (2). Here we found that GSIS is compromised in aging WT mice and is maintained in aging CgA-KO mice. Moreover, previous studies implicate age-related defects in post-receptor insulin signaling (2). While post-receptor insulin signaling was compromised in WT mice, insulin signaling remained intact in CgA-KO mice. Thus, it seems that the improved glucose tolerance, heightened insulin sensitivity and decreased hepatic glucose output are responsible for a healthier lifespan in CgA-KO mice.

### Inflammation and aging

One of the major and consistent changes that occurs during aging is the dysregulation of the immune response, leading to a chronic systemic inflammatory state (35). Healthy older people exhibit high serum concentrations of C-reactive protein (CRP) and other inflammatory mediators such as IL-6, IL-8, and TNF-α even in the absence of overt inflammatory disease (36). This so-called inflammaging has been reported to be associated with an increased risk of mortality in healthy and frail older adults (36), and increased levels of IL-6, CRP, and TNF-α, IL-1β and inflammasome-related genes are good predictors of all-cause mortality (36).

Conversely, lower levels of circulating inflammatory cytokines have been shown to correlate with good health outcomes, longevity, and reduced risk of death of older adults (37). The decreased levels of TNF-α, IFN-γ and CCL2 in older CgA-KO mice, therefore, further support the healthier lifespan in CgA-KO mice.

### Mitochondrial dysfunction and senescence

Aging is associated with mitochondrial dysfunction, including decreased oxidative capacity with increased oxidative damage, and hyperglycemia induced mitochondrial fragmentation/fission with enhanced respiration and increased ROS production. Our data found that aging WT mice had increased mitochondrial fission, which may result in increased ROS production (38). In contrast, aging CgA-KO mice showed more mitochondrial fusion, a process by which damaged mitochondria may acquire undamaged genetic material and maintain functionality (39). Moreover, electron transport has been reported to be optimal for ATP production in elongated/fused mitochondria (40). Therefore, the mitochondrial organization and the increased mitochondrial biogenesis in aging CgA-KO mice might support a healthy metabolism. In aging WT mice, the mitochondria are not only more fragmented, but their biogenesis is impaired owing to an age-associated decline in Pgc1a (41) and this is associated with a loss of mitochondrial content and function (41).

The 13 mtDNA-encoded proteins are all components of the respiratory chain or the ATP synthase, and oxidative phosphorylation subsides in the absence of mtDNA expression (42). The activity of complex I and IV decreases with age in the liver of mice (43). Consistent with this report, we found decreased expression of *MtNd1* and *MtAtp6* in aging WT mice, while the expression of those two genes remained consistently high in aging CgA-KO mice compared to WT mice.

The mtDNA has a higher rate of mutation (41) and has a higher rate of oxidative damage. There is substantial (up to 40%) reduction in rat liver mitochondria of old animals (24 months) in comparison with juvenile animals (3-4 months) (44). The current evidence suggests that mtDNA mutations and impaired OxPhos are primarily responsible for premature aging (45). Like the expression of *MtNd1* gene, nuclear encoded complex I genes *Ndufa9* and *Ndufs7* are also overexpressed in aging CgA-KO mice compared to aging WT mice. Together, these findings suggest that the organization and function of mitochondria are improved in aging CgA-KO mice.

### Hypertension

Age is a powerful risk factor for hypertension, death, and cardiovascular death ^83^. In the present study, we found a spontaneous reversal of hypertension in aging CgA-KO mice. Since supplementation of aging CgA-KO mice with the hyperglycemic and antidiabetic peptide PST increases both blood glucose and blood pressure, PST appears to be a key peptide whose inhibition or suppression may be responsible for a healthier lifespan.

## AUTHOR CONTRIBUTIONS

M.A.L., S.J., K.T. and S.K.M. researched data. M.A.L. and S.K.M. analyzed the data and wrote the manuscript. S.J., GvdB, and G.G. participated in discussion and reviewed/edited the manuscript. SKM conceived the study and made the graphics. M.A.L., S.J., K.T. and S.K.M. are the guarantor of this work and take responsibility for the integrity of the data and the accuracy of the data analysis.

## AVAILABILITY OF DATA AND MATERIAL

Primary data are available from the corresponding author on reasonable request.

## ACKNOWLEDGMENTS

Transmission Electron Microscopy was conducted at the Cellular & Molecular Medicine Electron Microscopy Core Facility at UCSD.

